# Genetic effect modification of cis-acting C-reactive protein variants in cardiometabolic disease status

**DOI:** 10.1101/2021.09.23.461369

**Authors:** Jie Zheng, Haotian Tang, Matthew Lyon, Neil M. Davies, Venexia Walker, James S. Floyd, Thomas R. Austin, Ali Shojaie, Bruce M. Psaty, Tom R. Gaunt, George Davey Smith

**Affiliations:** MRC Integrative Epidemiology Unit (IEU), Bristol Medical School, University of Bristol, Oakfield House, Oakfield Grove, Bristol, BS8 2BN, UK; Population Health Sciences, Bristol Medical School, University of Bristol, Barley House, Oakfield Grove, Bristol, BS8 2BN, United Kingdom; K.G. Jebsen Center for Genetic Epidemiology, Department of Public Health and Nursing, NTNU, Norwegian University of Science and Technology, Norway; Department of Surgery, University of Pennsylvania Perelman School of Medicine, Philadelphia, USA; Cardiovascular Health Research Unit, Departments of Medicine and Epidemiology and Medicine, University of Washington, Seattle, WA; Department of Biostatistics, University of Washington, University of Washington, Seattle, WA; Department of Health Services, University of Washington, Seattle, WA; NIHR Bristol Biomedical Research Centre, Bristol, UK

**Author notes:** Corresponding authorship. Equal first authorship (Jie Zheng, Haotian Tang). Equal last authorship (Tom R. Gaunt, George Davey Smith).

**Keywords:** cis-CRP variants, effect modification, disease status, body mass index

## Abstract

Mendelian randomization (MR) studies carried out among patients with a particular health condition should establish the genetic instrument influences the exposure in that subgroup, however this is normally investigated in the general population. Here, we investigated whether the genetic associations of four cis-acting C-reactive protein (CRP) variants differed between participants with and without three cardiometabolic conditions: obesity, type 2 diabetes, and cardiovascular disease. Associations of cis-genetic variants with CRP differed between obese and non-obese individuals. A multivariable analysis suggested strong independent associations of the gene-by-body mass index (BMI) interaction on CRP (P<1.18×10^−8^ for the CRP variants). Applying MR, we observed strong causal effect of BMI on CRP (P=2.14×10^−65^). In summary, our study indicates that genetic associations with CRP differ across disease sub-groups, with evidence to suggest that BMI is an effect modifier. MR studies of disease progression should report on the genetic instrument-exposure association in the disease subgroup under investigation.

## Introduction

Mendelian randomization (MR) is a genetic epidemiology method that can use genetic variants as instruments to estimate the causal role of a modifiable exposure (e.g. a protein) on an outcome (e.g. a disease) ^1^. Recent MR studies of molecular phenotypes showed that it can provide reliable evidence of potential drug target efficacy ^2^. In such studies the causes of disease onset are usually being investigated. In order to study the effects of potential therapeutic interventions on disease, MR is applied to the progression of disease once it has developed ^3,4^.

Till now, most of the existing MR studies assume that the genetic effects of instruments are consistent in different subgroups of the population, including patients with particular conditions. However, naive use of genetic instruments generated in a general population as proxy instruments in disease subgroups may yield biased estimates of causal relationships between the exposure and outcome, especially for MR of disease progression. For example, C-reactive protein (CRP) is a biomarker of inflammation associated with atherosclerosis, and atherosclerosis may modify the association of genetic instruments with CRP. Previous observational studies have also suggested that higher BMI/obesity is associated with higher CRP level ^5,6^, where previous MR study further showed the causal relationship of BMI on CRP ^7^. Thus, people with cardiometabolic disorders may have different genetic variant associations with CRP than those without these disorders, and MR will be biased if these differences are not accounted for. Addressing differences in genetic effects across subgroups may therefore be important when using MR to predict drug effects on disease progression.

In this study, we investigated whether the associations of genetic variants with CRP differed between people with and without cardiometabolic disease. We conducted a set of statistical analyses to investigate three scientific questions related to genetic effect modification in disease status: i) do the genetic associations of the cis-CRP variants differ between those with and without cardiometabolic disease; ii) does gene-by-environment (G×E) interaction play a role in influencing CRP level; iii) does genetically predicted BMI contribute to the genetic effect modification of CRP. Overall, our analysis provides a step towards better understanding of genetic effect modification of molecular phenotypes in disease status.

## Results

### Study characters

We selected four well-characterised cis-CRP variants (rs3093077, rs1205, rs1130864 and rs1800947) from a previous MR study of CRP ^8^. The individual level CRP measurements, genotypes for the four cis-CRP variants, and cardiometabolic disease status were obtained from UK Biobank. The sample size of each disease subgroups after sample selection is demonstrated in **Supplementary Figure 1**. More details of sample selection could be found in Methods.

### Estimation of genetic effect modification of CRP variants in cardiometabolic disease status

First, we investigated if genetic associations with CRP differed in participants with and without cardiometabolic diseases, which we estimated the genetic association of CRP (in standard deviation [SD] unit) of each cis-CRP variants stratified by obesity status, diabetes status and cardiovascular disease (CVD) status. As shown in **Figure 1**, we found strong evidence of heterogeneity in the SNP associations for all four cis-CRP variants in the obese (BMI≥30), non-obese (BMI<30) and overall samples. The SNPs were more strongly associated with CRP among obese participants than non-obese (**Supplementary Table 1**). We found similar heterogeneity in participants with and without diabetes, but these estimates were less precise. In contrast, we found little evidence of heterogeneity by CVD status (**Supplementary Table 1**). Given the heterogeneity in the SNP-CRP associations for the obesity and diabetes subgroups, we further investigated these two diseases in a secondary analysis.

**Figure 1.**
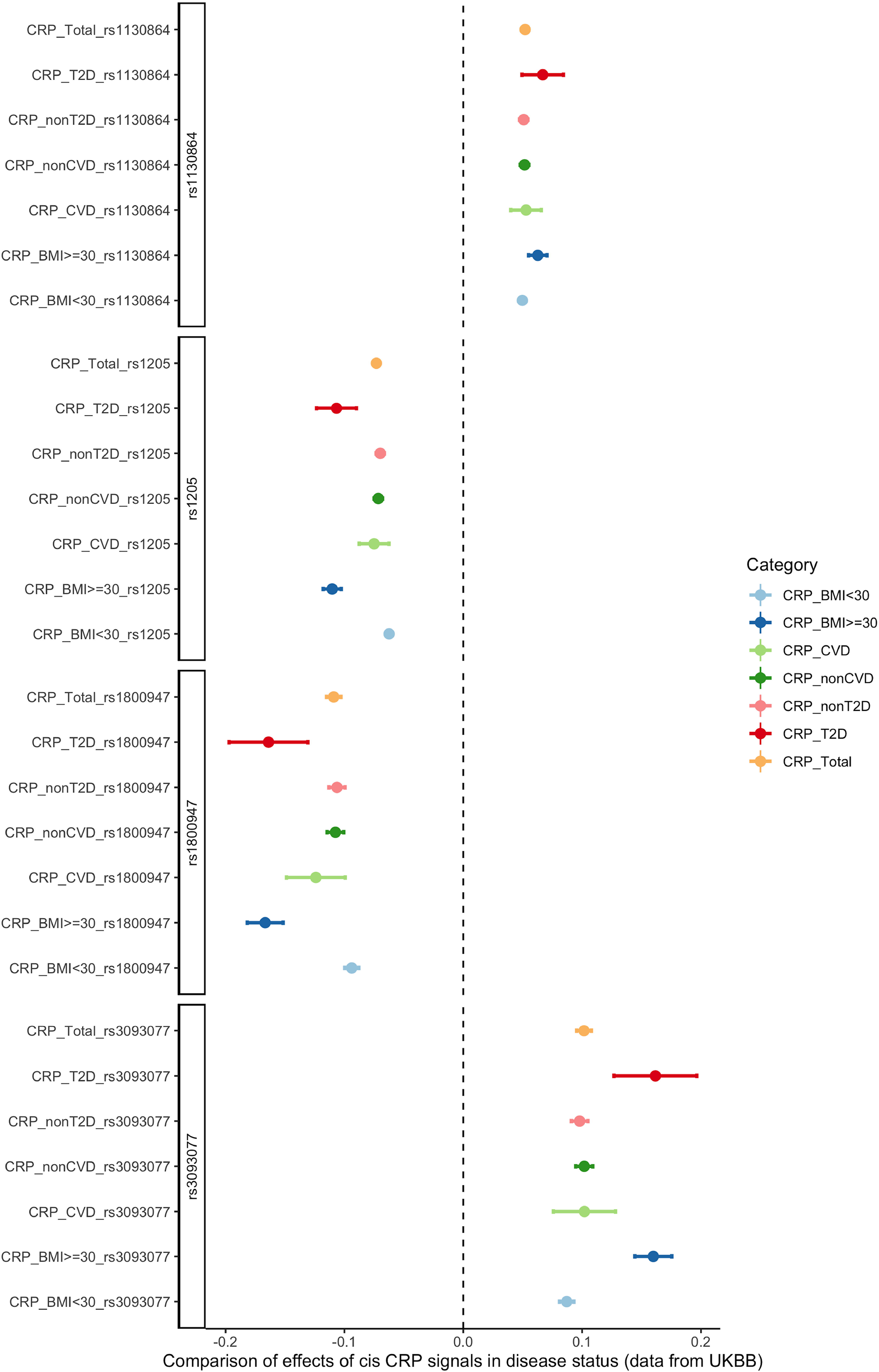
Genetic effect of four cis-CRP variants in disease status. Seven groups of samples were tested: overall (orange), obese and non-obese (dark and light red), diabetes and non-diabetes (dark and light green), CVD and non-CVD (dark and light blue).

### Estimation of the influences of gene-by-environmental interaction on CRP

Second, we investigated whether gene-by-environment (G×E) interactions influenced CRP level, which gives evidence on whether the SNP effect is modified by an environmental factor (e.g. BMI).

**Figure** *2* illustrates the independent G×BMI effects of the four cis-CRP variants on CRP. We found evidence of G×BMI interactions for all four cis-CRP variants (P<1.18×10^−8^), with the main SNP effects remaining strong (P<4.14×10^−242^, **Supplementary Table 2**). In addition, we found evidence of GxHbA1c interaction for three of four CRP variants, where rs3093077, rs1205 and rs1800947 showed strong evidence of an interaction with HbA1c (P<3.14×10^−8^), but rs1130864 had weaker evidence (P=0.03) (**Supplementary Figure 2)**. There was little evidence that including an interaction with HbA1c attenuated the association of the SNPs with CRP (**Supplementary Table 3**). Given the potential influence of scale on the interaction model, we ran the international analysis using exposure (or) outcome on the additive and multiplicative scale (e.g. took the log or exponential for the exposure or outcome and run the same interaction analysis). A robust interaction term was seen with both models G×BMI (results not shown). As we observed strong G×BMI effects for all four CRP variants, we followed up this finding in the third analysis.

**Figure 2.**
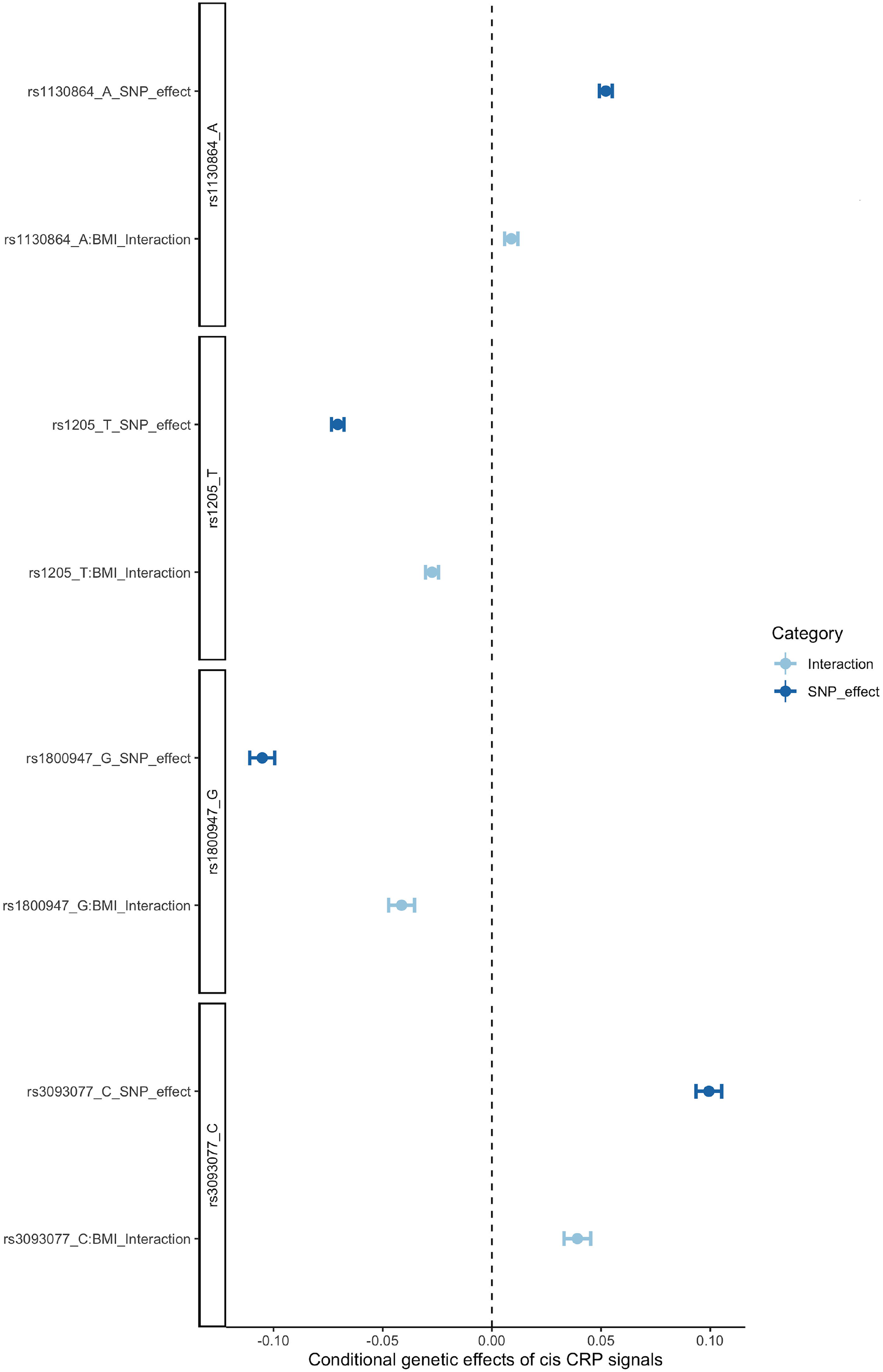
Genetic and gene by BMI interaction effects of four cis-CRP variants.

### Estimation of the contribution of body mass index on CRP genetic effect modification

Third, we investigated whether BMI contributes to the modification of genetic association of CRP. We performed a genetic association analysis of CRP stratified by the genetic score of BMI (gsBMI). The gsBMI was constructed using 73 BMI associated SNPs from a previous paper ^9^ (**Supplementary Table 4**). For the stratification of gsBMI, we estimated gsBMI-CRP associations within five quintiles of the gsBMI (20% quantile each group)..

**Figure** *3* shows the genetic effects of cis-CRP variants stratified into gsBMI quintiles. We found evidence of a linear trend suggests that as the genetic score of BMI increased, the genetic association of rs3093077, rs1205 and rs1800947 with CRP also increased (**Supplementary Figure 3** and **Supplementary Table 5**). This suggests that BMI likely modifies the genetic associations of these three variants with CRP. We further conducted a one-sample MR to estimate the causal influence of BMI on CRP, where the gsBMI of the 73 BMI SNPs ^9^ (**Supplementary Table 4**) were used as weight to construct the exposure. The one-sample MR suggested a robust positive causal effect of BMI on CRP (β=0.198, beta refers to SD unit change of CRP per SD unit change of BMI; 95%CI=0.175 to 0.220; P=2.14×10^−65^; **Figure 4** and **Supplementary Table 6**). This further confirmed the causal role of BMI on CRP.

**Figure 3.**
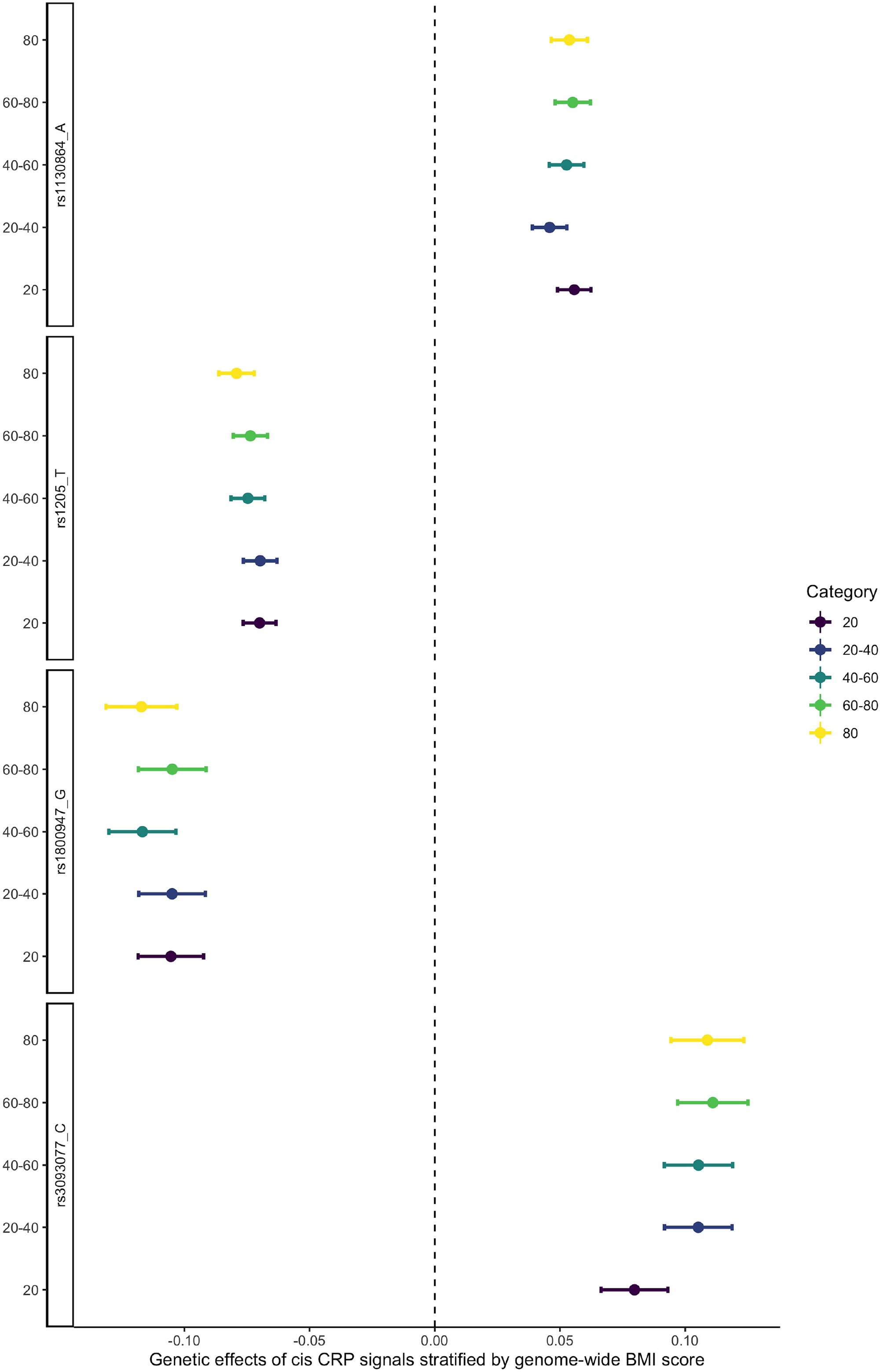
Genetic effect of four cis-CRP variants stratified by BMI genetic score. The results were stratified into five categories: BMI score below 20% quantile (dark purple); from 20% to 40% quantile (dark blue); 40% to 60% quantile (light blue); 60% to 80% quantile (light green) and over 80% quantile (yellow).

**Figure 4.**
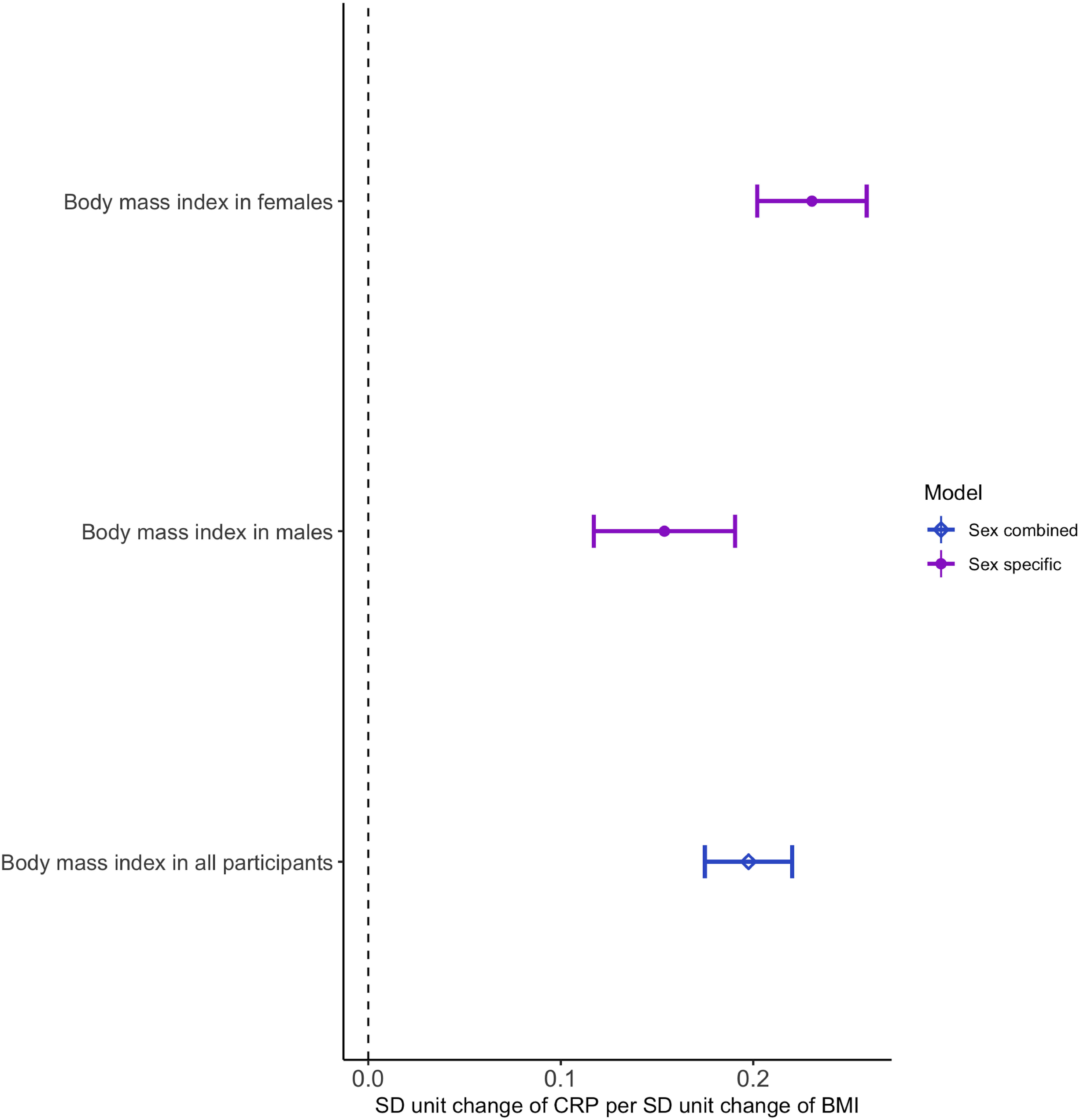
One-sample Mendelian randomization estimates of body mass index on C-reactive protein. The results were stratified into two models: sex specific Mendelian randomization analysis in Females and males; sex combined Mendelian randomization analysis in all participants.

### Estimation of sex-specific effect in genetic effect modification

Finally, we investigated whether sex modified genetic associations in sensitivity analyses. We estimated the heterogeneity of the genetic association with CRP stratified by obese and non-obese status for males and females separately. In general, the associations of the SNPs with CRP were larger in females, although these differences were not statistically robust (**Supplementary Figure 4** and **Supplementary Table 7A**). We found strong evidence of an interaction between BMI and all genetic variants in both males and females (**Supplementary Figure 5** and **Supplementary Table 7B**). For the analysis stratified by genetic score of BMI, we observed a similar linear trend for rs3093077, rs1205 and rs1800947, but found an unclear trend for rs1130864 (**Supplementary Figure 6** and **Supplementary Table 7C**). The sex-stratified one-sample MR further confirmed the causal role of BMI on CRP in males and females (**Figure 4** and **Supplementary Table 6**). In general, the sex specific analyses suggested a similar trend of genetic effect modification of CRP in males and females, with slightly larger effect estimates in females.

## Discussion

MR studies investigating drug target effects (such as level of a protein) on progression of a disease have high potential value in predicting therapeutic drug trial efficacy ^4^. In this study, we investigated the genetic effect modification in disease status using protein level of CRP as a case. We observed strong evidence to support that genetic associations with CRP differ between obese/non-obese and diabetic/non-diabetic individuals. The interaction analysis showed that gene by BMI and gene by HbA1c interaction showed independent associations with CRP. Further genetic association and MR analyses provided strong evidence to suggest that BMI is an effect modifier on CRP. These findings highlight the importance of considering effect modification by disease status in future MR studies of disease progression.

Our study has several implications. First, CRP is unlikely to cause the disease subgroups we investigated ^8^. Therefore collider bias, which is also known as selection bias or sampling bias ^10^, is unlikely to have impacts for this study. However, future MR studies in the disease progression setting, for example IL6R on CVD ^11^, may be affected and collider bias should then be addressed using methods such as the Slope-Hunter approach^12^.

Second, we estimated the G×E effect and provided strong evidence of an independent effect of G×BMI on CRP. Of note, the interaction effect is model dependent. With increased sample size of biobanks, we are likely to identify an “interaction” for exposure with a main effect on a certain scale. To avoid misuse of interaction ^13^, we highly recommend estimating the interaction on different scales of the exposure/outcome. As a demonstration, we applied the interaction model on both the additive and multiplicative scales in this study.

Third, our study is a hypothesis-driven study focusing on whether genetic associations with CRP differ across three common cardiometabolic diseases. A recent MR study has provided evidence for a widespread influence of BMI on the human proteome ^14^. Our MR results suggested that increasing BMI level is associated with increased CRP level, which confirmed the finding from a previous MR study using an FTO SNP as instrument ^7^. Our study provides additional evidence of this, suggesting that BMI inflates genetic associations with CRP in obese patients compared to non-obese individuals. In future studies, causal effects for promising putative drug targets may differ by disease status, and the stratified analyses proposed in this study could be more widely applied.

Finally, we focus on just one protein (CRP), which has been widely analysed in population studies, enabling us to obtain individual level CRP data from over 360,000 individuals. For most of the remaining proteins, the sample sizes for the genetic studies are still limited ^15–17,18^ and may not provide sufficient statistical power for this type of analysis. However, we anticipate that we will be able to systematically estimate the influence of disease status using MR of plasma proteins in roughly 50,000 samples from the proteome resources in the UK Biobank when these data are released.

In conclusion, our study demonstrates that genetic associations with CRP differs between obese and non-obese individuals. This heterogeneity may influence MR results. Therefore, it is important to obtain associations of genetic variants with exposure from comparable samples to those being used to instrument disease-status in future MR studies of disease progression.

## Methods

### Study samples

The UK Biobank is a large-scale biomedical database and research resource, containing in-depth genetic and health information from half a million UK participants. For this study, we obtained individual level CRP measurements, genotype data of the CRP variants (more details in Genetic instrument selection section), and cardiometabolic disease status from UK Biobank. We evaluated three primary cardiometabolic conditions: obesity (BMI≥30), type 2 diabetes (ICD10 code E11), and cardiovascular disease (CVD, ICD10 codes I21-I25). We also evaluated two continuous cardiometabolic measures: BMI (UKBB data field 23104), and haemoglobin A1c (HbA1c, UKBB data field 30750). UK Biobank received ethical approval from the North West Multi-centre Research Ethics Committee (MREC reference for UK Biobank is 11/NW/0382).

### Sample selection and quality control

We restricted statistical analysis to 365,149 unrelated European samples to control for the influence of population structure and relatedness (**Supplementary Figure 1**). We removed outliers outside four standard deviation (SD) units from the mean for the three quantitative phenotypes, CRP, BMI and HbA1c, to reduce bias in the association estimate. For all statistical analyses, we included age, sex, genotyping array (noted as chip) and the first 40 principal components as covariates in the statistical model to control for the influence of these variables and population structure on the association estimation. The sample size for each disease subgroup after selection is demonstrated in **Supplementary Figure 1**.

### Genetic instrument selection

For the instrument selection, we aimed to identify genetic variants that robustly associated with CRP and showed the most promising ability in genetic and MR analysis. In common practice of proteome MR analysis, cis-acting variants, defined as genetic variants within or nearby the protein coding gene, are more likely to have protein specific effects than trans-acting variants ^15^. They have been used as genetic instruments for much of the published drug target MR ^11,19^. Here, we selected four well-characterised cis-CRP variants (rs3093077, rs1205, rs1130864 and rs1800947) from a previous MR study of CRP ^8^. These variants were used for the genetic association and MR analyses.

### Statistical analyses

We conducted three sets of statistical analyses to investigate the genetic effect modification in disease status for the cis-CRP variants: i) estimating the genetic effect modification of the four cis-CRP variants in those with and without the three types of cardiometabolic diseases: obesity, type 2 diabetes and cardiovascular disease (CVD); ii) estimating the independent effect of gene-by-environment (G×E) interaction on CRP; iii) estimating the influence of BMI on genetic effect modification of CRP in disease status.

#### Genetic association analysis stratified by disease status

For the first analysis, we applied a linear regression to estimate the association of CRP (in SD unit) and the genotype of each cis-CRP variant stratified by obesity status, diabetes status, and CVD status. As a reference, we also estimated the genetic associations of the cis-CRP variants in the whole UK Biobank sample. We draw forest plots to compare the genetic effect changes in each subgroup of patients.

#### Gene-by-environment interaction effect on CRP

For the second analysis, we estimated the effect of gene-by-environment interaction on CRP using the following multivariable regression model including a G×E interaction term:

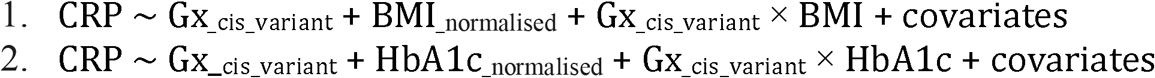

Where Gx is the genotype of one of the cis-CRP variants. BMI and HbA1c levels (in SD units) were normalised using their population median to allow consistent directionality for the main SNP effect and the interaction effect.

Given the scale of the exposure and outcome may influence the effect of the interaction on the outcomes, we therefore conducted a sensitivity analysis to estimate such potential influence. We ran the interaction analysis using the exposure (or outcome) on both the additive and multiplicative scale. For example, we took the log (or exponential) for the exposure or outcome and run the interaction model to test whether the interaction effect on CRP still exist.

#### Genetic association analysis of CRP stratified by genetic score of BMI

For the third analysis, we performed a linear regression model of CRP (in SD unit) against the genotype of each cis-CRP variant stratified by the genetic score of BMI (gsBMI). The gsBMI was constructed using 73 BMI associated SNPs that have been widely used in previous MR studies ^9^ (**Supplementary Table 4**). To further explore the linearity of the relationships, we estimated the gsBMI-CRP association within five subgroups divided by the quintile of quintiles of the gsBMI (20% quantiles each subgroup). The forest plot was created to visualise the linearity.

#### Mendelian randomization analysis of BMI on CRP

For the third analysis, we further conducted a one-sample MR to estimate the causal influence of BMI on CRP. The gsBMI of the same 73 BMI SNPs ^9^ (**Supplementary Table 4**) were used as weight to construct the exposure. The CRP level in UK Biobank were used as outcome for the MR. The two-stage least square approach were applied to estimate the causal effect of BMI on CRP.

#### Sex-specific analysis

Finally, given the potential influence of different BMI in males and females, we estimated the heterogeneity of the genetic association with CRP stratified by obese and non-obese status for males and females separately. The same genetic association and MR analyses were applied for this sensitivity analysis.

All statistical analyses were conducted using R v4.0.2.

## Supplemental information

Supplemental Tables and Figures can be found online.

## References

1. Davey Smith, G. & Ebrahim, S. ‘Mendelian randomization’: can genetic epidemiology contribute to understanding environmental determinants of disease? Int. J. Epidemiol. 32, 1–22 (2003).

2. Zheng, J. et al. Phenome-wide Mendelian randomization mapping the influence of the plasma proteome on complex diseases. Nat. Genet. 52, 1122–1131 (2020).

3. Davey Smith, G., Paternoster, L. & Relton, C. When Will Mendelian Randomization Become Relevant for Clinical Practice and Public Health? JAMA 317, 589–591 (2017).

4. Paternoster, L., Tilling, K. M. & Smith, G. D. Genetic Epidemiology And Mendelian Randomization For Informing Disease Therapeutics: Conceptual And Methodological Challenges. (2017) doi:10.1101/126599.

5. Visser, M., Bouter, L. M., McQuillan, G. M., Wener, M. H. & Harris, T. B. Elevated C-reactive protein levels in overweight and obese adults. JAMA 282, 2131–2135 (1999).

6. Aronson, D. et al. Obesity is the major determinant of elevated C-reactive protein in subjects with the metabolic syndrome. Int. J. Obes. Relat. Metab. Disord. 28, 674–679 (2004).

7. Timpson, N. J. et al. C-reactive protein levels and body mass index: elucidating direction of causation through reciprocal Mendelian randomization. Int. J. Obes. 35, 300–308 (2011).

8. C Reactive Protein Coronary Heart Disease Genetics Collaboration (CCGC) et al. Association between C reactive protein and coronary heart disease: mendelian randomisation analysis based on individual participant data. BMJ 342, d548 (2011).

9. Sun, Y.-Q. et al. Body mass index and all cause mortality in HUNT and UK Biobank studies: linear and non-linear mendelian randomisation analyses. BMJ 364, l1042 (2019).

10. Munafò, M. R., Tilling, K., Taylor, A. E., Evans, D. M. & Davey Smith, G. Collider scope: when selection bias can substantially influence observed associations. Int. J. Epidemiol. (2017) doi:10.1093/ije/dyx206.

11. Interleukin-6 Receptor Mendelian Randomisation Analysis (IL6R MR) Consortium et al. The interleukin-6 receptor as a target for prevention of coronary heart disease: a mendelian randomisation analysis. Lancet 379, 1214–1224 (2012).

12. Mahmoud, O., Dudbridge, F., Smith, G. D., Munafo, M. & Tilling, K. Slope-Hunter: A robust method for index-event bias correction in genome-wide association studies of subsequent traits. (2020) doi:10.1101/2020.01.31.928077.

13. Davey Smith, G. Post-Modern Epidemiology: When Methods Meet Matter. Am. J. Epidemiol. 188, 1410–1419 (2019).

14. Goudswaard, L. J. et al. Effects of adiposity on the human plasma proteome: observational and Mendelian randomisation estimates. Int. J. Obes. (2021) doi:10.1038/s41366-021-00896-1.

15. Sun, B. B. et al. Genomic atlas of the human plasma proteome. Nature 558, 73–79 (2018).

16. Folkersen, L. et al. Mapping of 79 loci for 83 plasma protein biomarkers in cardiovascular disease. PLoS Genet. 13, e1006706 (2017).

17. Emilsson, V. et al. Co-regulatory networks of human serum proteins link genetics to disease. Science (2018) doi:10.1126/science.aaq1327.

18. Yao, C. et al. Genome-wide mapping of plasma protein QTLs identifies putatively causal genes and pathways for cardiovascular disease. Nat. Commun. 9, 3268 (2018).

19. Holmes, M. V., Richardson, T. G., Ference, B. A., Davies, N. M. & Davey Smith, G. Integrating genomics with biomarkers and therapeutic targets to invigorate cardiovascular drug development. Nat. Rev. Cardiol. 18, 435–453 (2021).

