## Supplementary Figure for "Genetic effect modification of cis-acting C-reactive protein variants in cardiometabolic disease status"

### Supplementary Figures

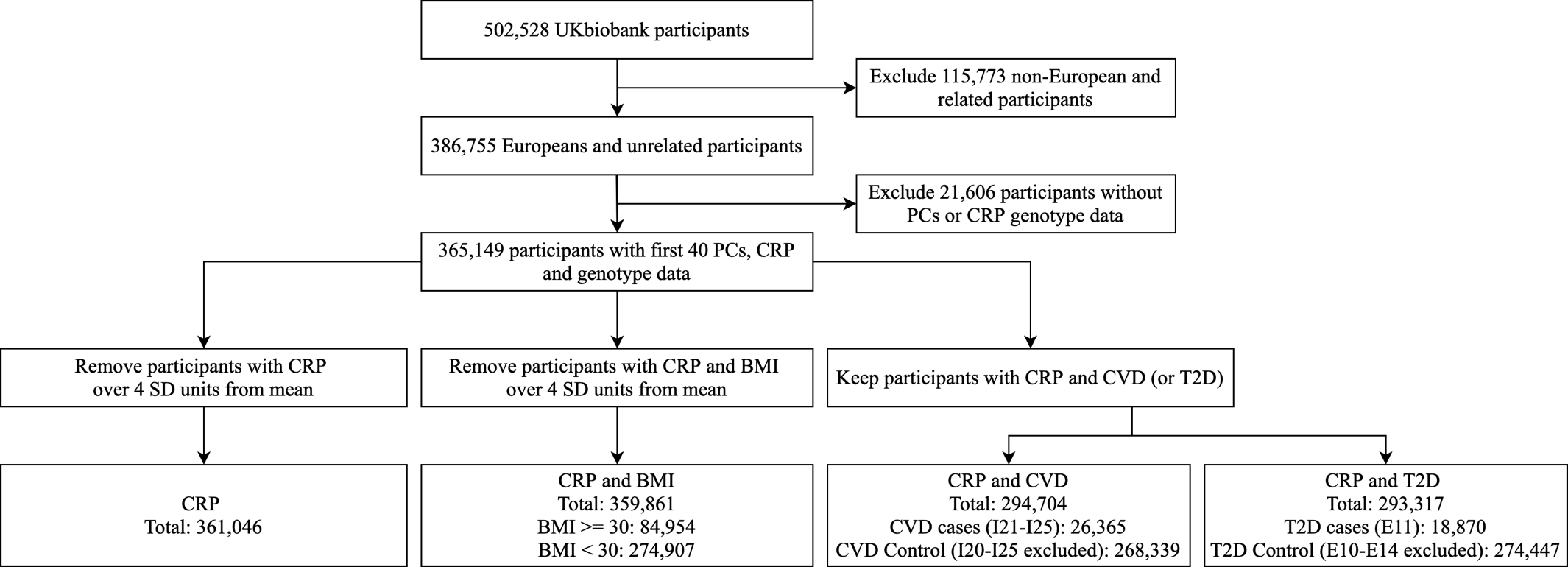

Supplementary Figure 1. Flowchart of sample selection of the study. Notation: PCs: principal components; SD: standard deviation; CRP: C-reactive protein; BMI: body mass index; CVD: cardiovascular disease; T2D: type 2 diabetes.

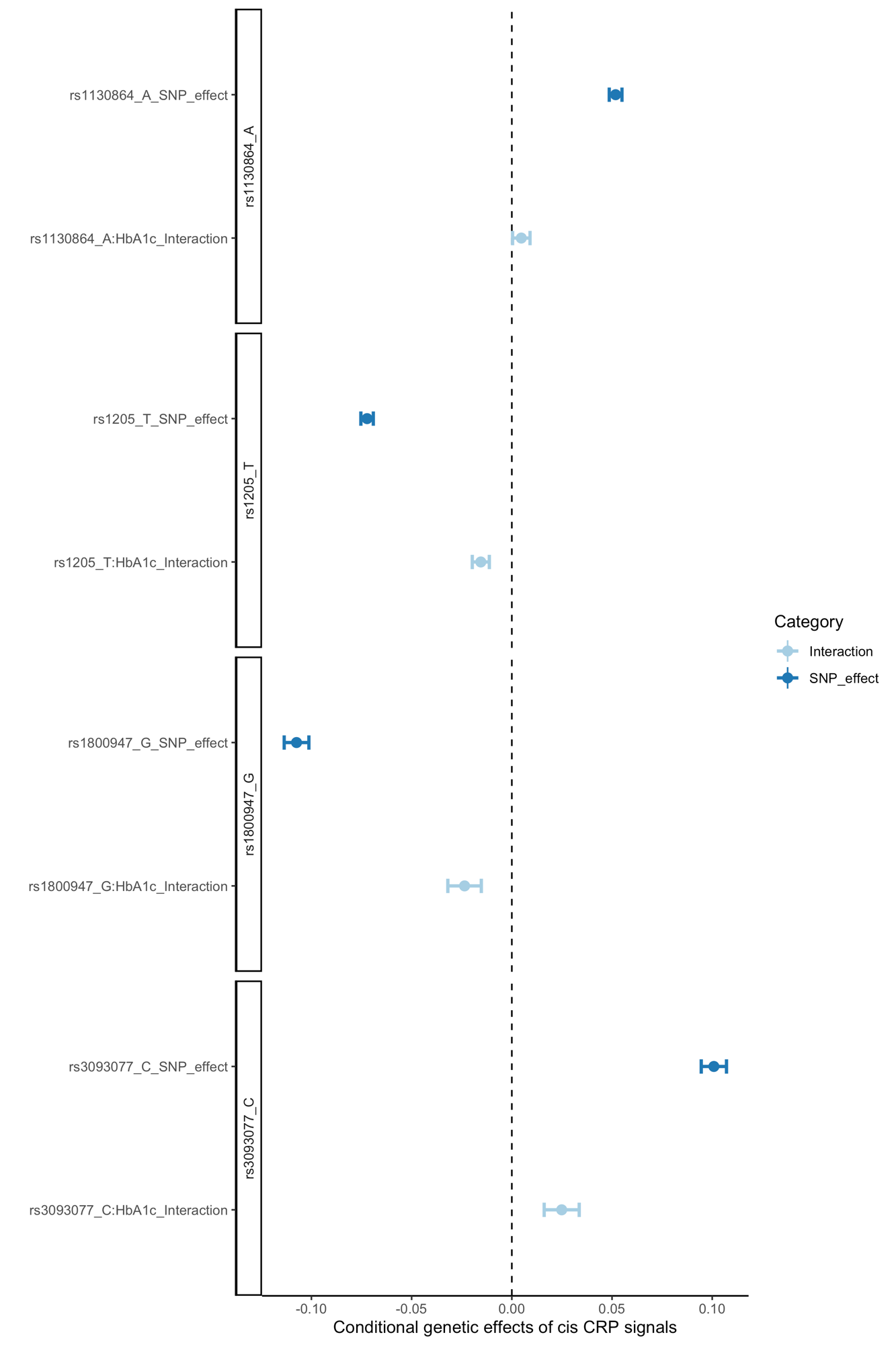

Supplementary Figure 2. Genetic and gene by HbA1c interaction effects of four cis-CRP variants.

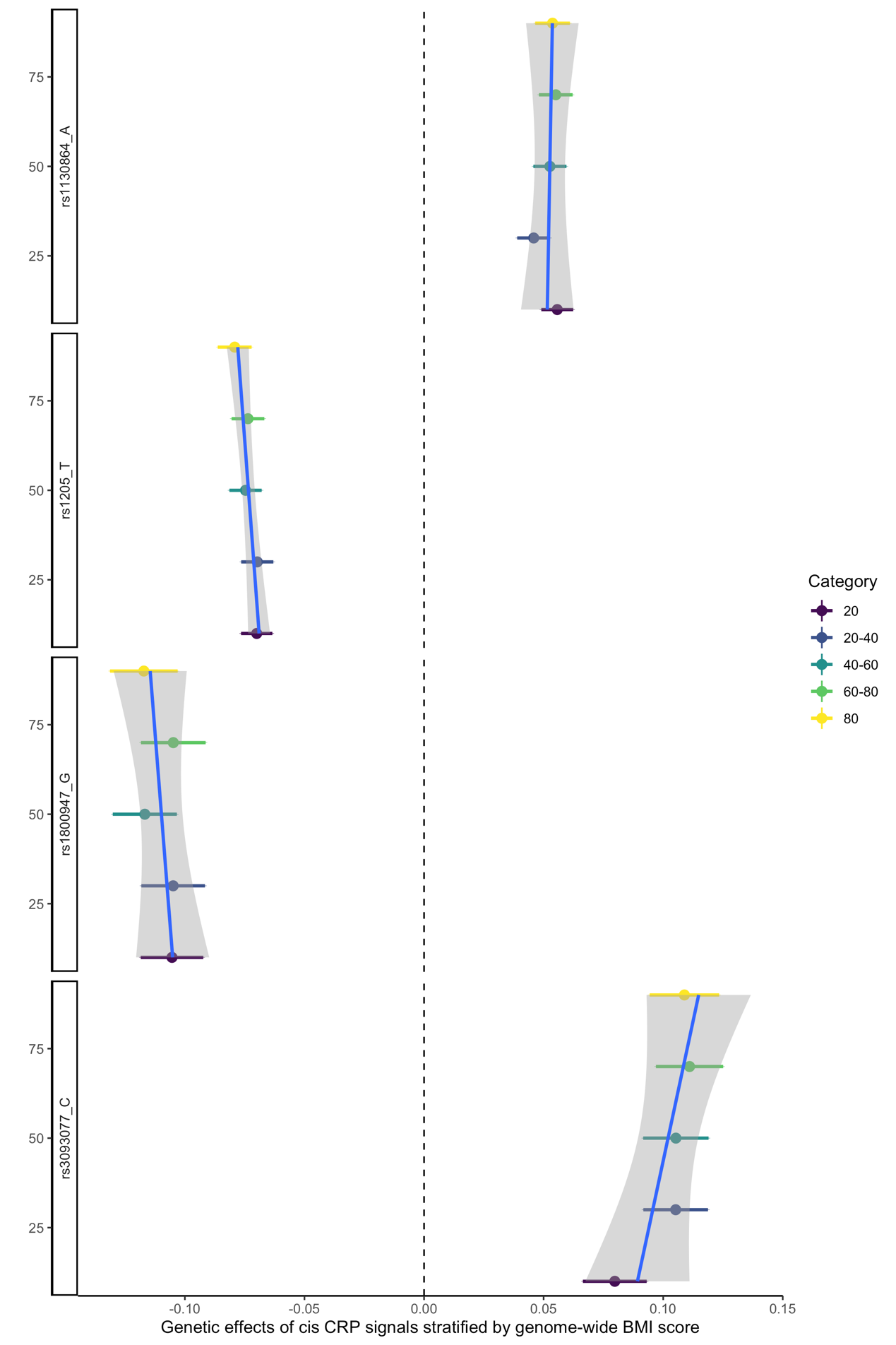

Supplementary Figure 3. Genetic effect of four cis-CRP variants stratified by BMI genetic score with linear trend line. The results were stratified into five categories: BMI score below 20% quantile (dark purple); from 20% to 40% quantile (dark blue); 40% to 60% quantile (light blue); 60% to 80% quantile (light green) and over 80% quantile (yellow). Linear trend line was in blue. The 95% confidence interval of the linear trend line was in grey.

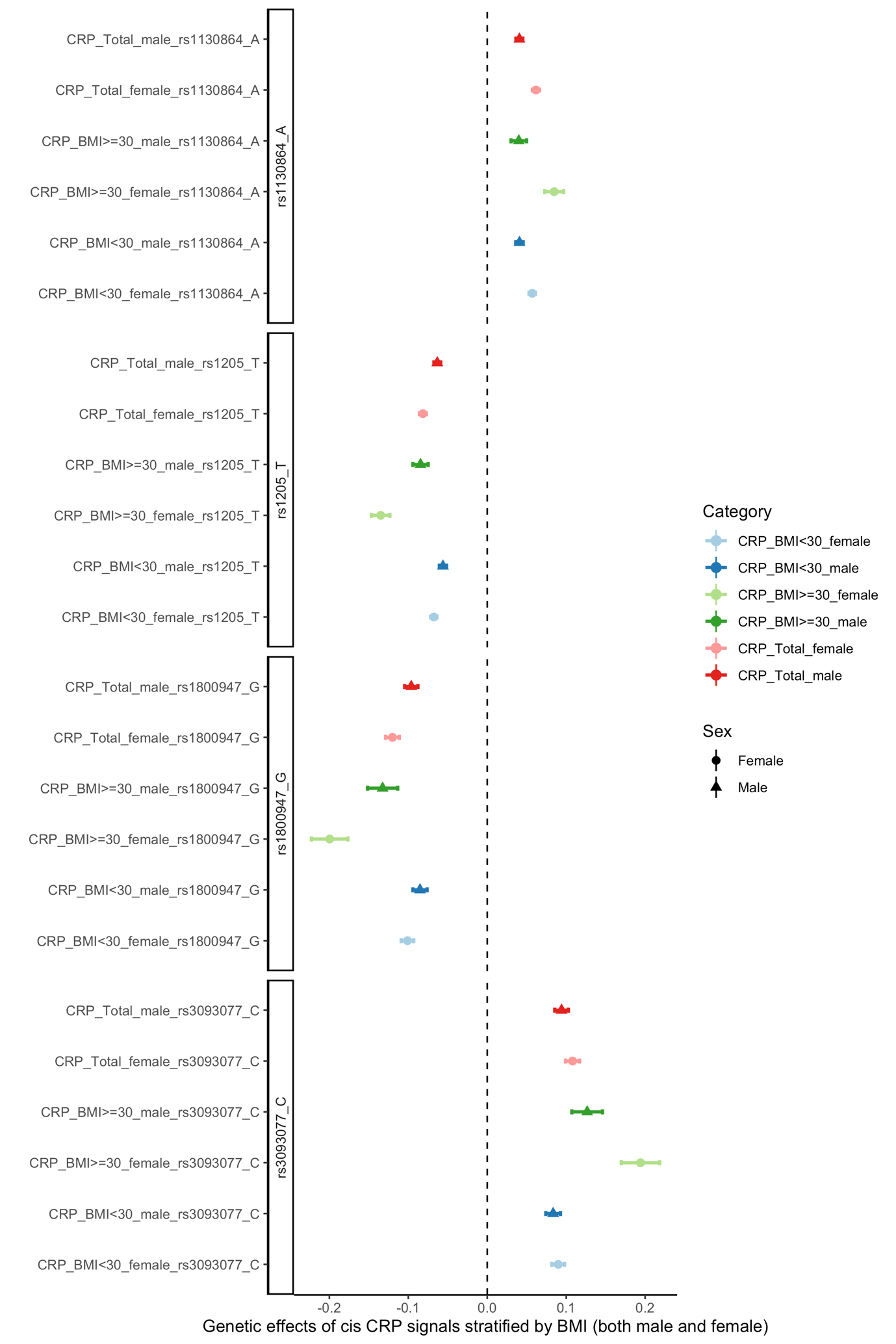

Supplementary Figure 4. Male and female only genetic effects stratified by BMI. Circle dot refers to female effect, triangle dot refers to male effect.

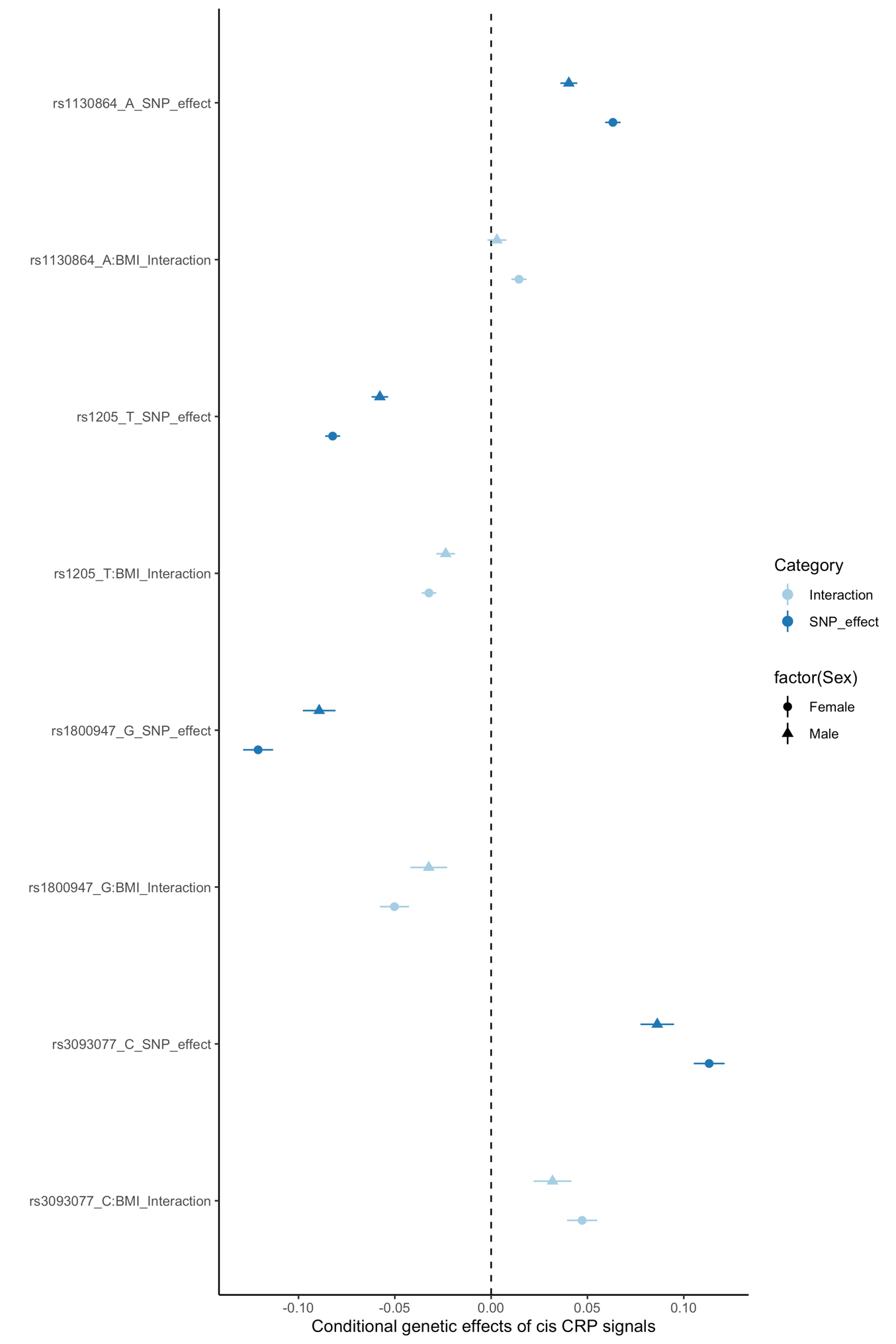

Supplementary Figure 5. Male and female only gene by BMI effects of the four cis-CRP variants*.* Circle dot refers to female effect, triangle dot refers to male effect.

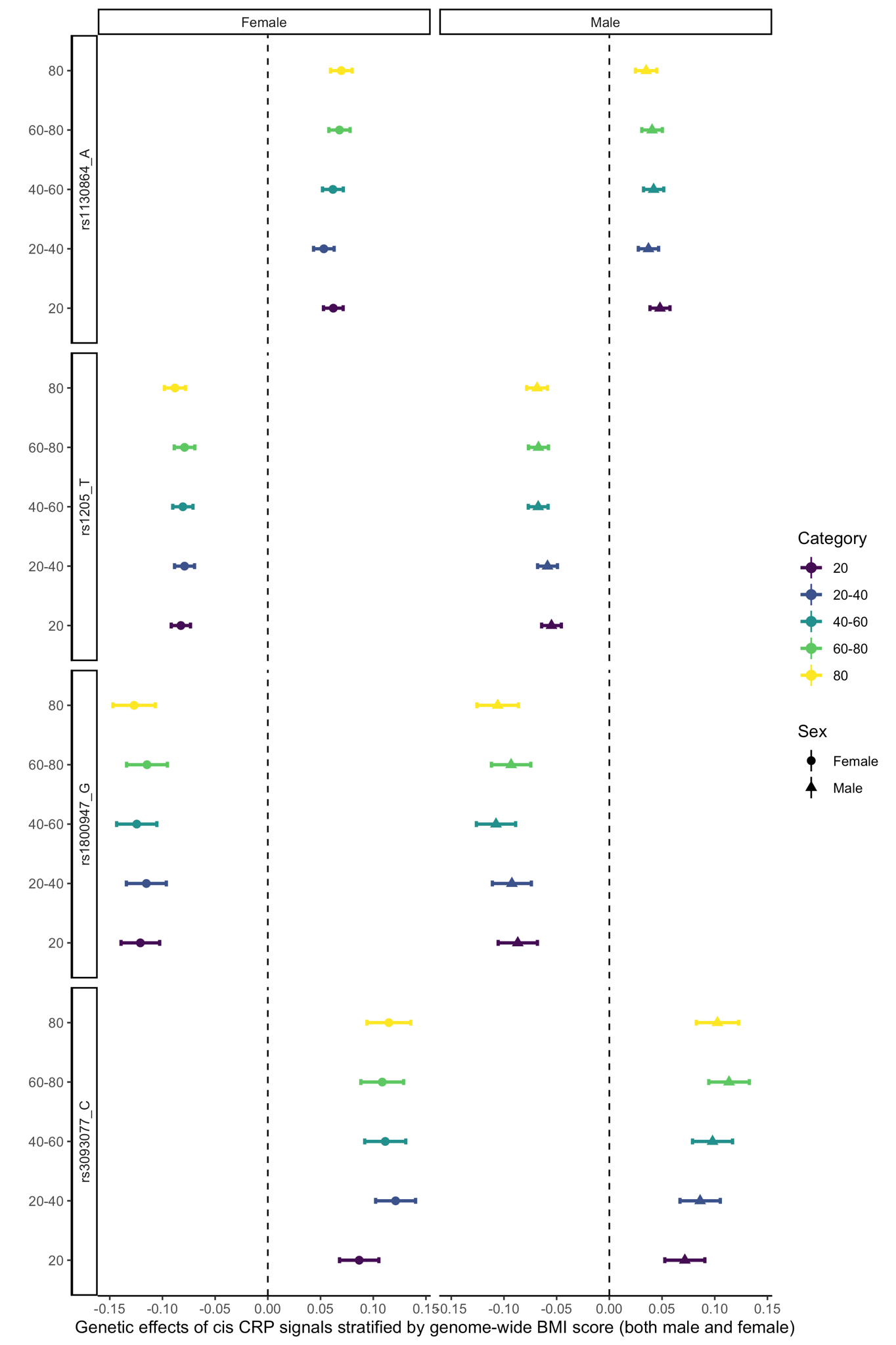

Supplementary Figure 6. Male and female only genetic effects stratified by BMI score*.* Circle dot refers to female effect, triangle dot refers to male effect.
